# Directed Evolution of Acoustic Reporter Genes Using High-Throughput Acoustic Screening

**DOI:** 10.1101/2024.03.30.587094

**Authors:** Robert C. Hurt, Zhiyang Jin, Mohamed Soufi, Katie K. Wong, Daniel P. Sawyer, Hao K. Shen, Przemysław Dutka, Ramya Deshpande, Ruby Zhang, David R. Mittelstein, Mikhail G. Shapiro

## Abstract

A major challenge in the fields of biological imaging and synthetic biology is noninvasively visualizing the functions of natural and engineered cells inside opaque samples such as living animals. One promising technology that addresses this limitation is ultrasound (US), with its penetration depth of several cm and spatial resolution on the order of 100 µm.^1^ Within the past decade, reporter genes for US have been introduced^2,3^ and engineered^4,5^ to link cellular functions to US signals via heterologous expression in commensal bacteria and mammalian cells. These acoustic reporter genes (ARGs) represent a novel class of genetically encoded US contrast agent, and are based on air-filled protein nanostructures called gas vesicles (GVs).^6^ Just as the discovery of fluorescent proteins was followed by the improvement and diversification of their optical properties through directed evolution, here we describe the evolution of GVs as acoustic reporters. To accomplish this task, we establish high-throughput, semi-automated acoustic screening of ARGs in bacterial cultures and use it to screen mutant libraries for variants with increased nonlinear US scattering. Starting with scanning site saturation libraries for two homologs of the primary GV structural protein, GvpA/B, two rounds of evolution resulted in GV variants with 5- and 14-fold stronger acoustic signals than the parent proteins. We anticipate that this and similar approaches will help high-throughput protein engineering play as large a role in the development of acoustic biomolecules as it has for their fluorescent counterparts.

## INTRODUCTION

Acoustic reporter genes (ARGs)—genetically encoded reporters that enable the imaging of gene expression using ultrasound (US)—were first introduced to bacteria in 2018^2^ and subsequently to mammalian cells in 2019.^3^ ARGs are based on genetically encoded, gas-filled protein nanostructures called gas vesicles (GVs) that originally evolved in buoyant microbes.^6,7^ GVs scatter US due to the difference in the density and compressibility of their gaseous interior relative to a surrounding aqueous medium.^8^ GVs have been the subject of intense study^7–14^, development,^4^ and application^5,15–23^ in recent years.^1,24,25^ ARGs have received considerable attention due to their ability to enable noninvasive, long-term, real-time imaging of gene expression in both bacterial and mammalian cells deep inside living organisms: in particular, ARGs have been used to image tumor growth^3,4^ and colonization by therapeutic bacteria,^4^ protease activity,^5^ phagolysosomal function,^9^ and intracellular Ca^2+^ dynamics.^10^ However, despite several successful efforts to engineer the acoustic and expression properties of ARGs, further improvements to the performance of ARGs are needed to enable their most impactful applications.

Unfortunately, the methods currently available for ARG engineering and acoustic characterization are low-throughput, complex to implement, and require a great deal of hands-on time per sample. In particular, manual loading and imaging of individual samples limits throughput to a handful of samples per day. In contrast, the state-of-the-art high-throughput methods used to engineer fluorescent proteins can process far larger libraries in shorter times, with less intervention from users: plate readers can assay thousands of samples per run, and flow cytometers have been used to screen libraries of 10^8^ mutants in a single experiment.^11^ In the past few decades, a growing suite of protein engineering techniques have been developed^12^ and applied with remarkable success to improving fluorescent proteins, opsins, Cas proteins, and other biotechnology tools, but these methods often require the screening of libraries containing thousands of members or more.^13^ Thus, the low throughput of current acoustic screening methods prevents the effective use of most of the tools needed to unlock the full potential of ARGs.

In this study, we developed a high-throughput, semi-automated pipeline for acoustic screening of ARGs, and used it to evolve two ARG clusters to improve their nonlinear acoustic signals. Our acoustic plate reader (APR) system is capable of collecting acoustic data on up to 1152 ARG samples in a single automated scan, and includes graphical user interfaces (GUIs) for data collection and processing. The APR workflow facilitates faster, more reliable, and more standardized acoustic screening of ARG samples, requiring significantly less hands-on time than current methods. Using this pipeline, we improved the nonlinear acoustic signal produced by two ARG clusters—derived from *Anabaena flos-aquae* and *Bacillus megaterium*—by 5- and 14-fold, respectively, when expressed at physiological temperature. Microscopy revealed that these evolved ARG clusters produce more GVs per cell than their parents.

## RESULTS

### A high-throughput directed evolution workflow for ARGs

GVs are known to respond to US in three regimes, depending on the input pressure applied: linear scattering, nonlinear scattering, and collapse^14,15^ (Figure 1A). Of particular interest for *in vivo* imaging is the nonlinear scattering regime in which GVs produce significantly more contrast than tissue, putatively by “buckling” of their shells.^14–18^ This effect has been exploited previously to non-destructively image GV-expressing bacterial and mammalian cells *in vivo* with high specificity,^4^ and enhancing this nonlinear US scattering phenotype is a top priority of current ARG engineering efforts.

**Figure 1.**
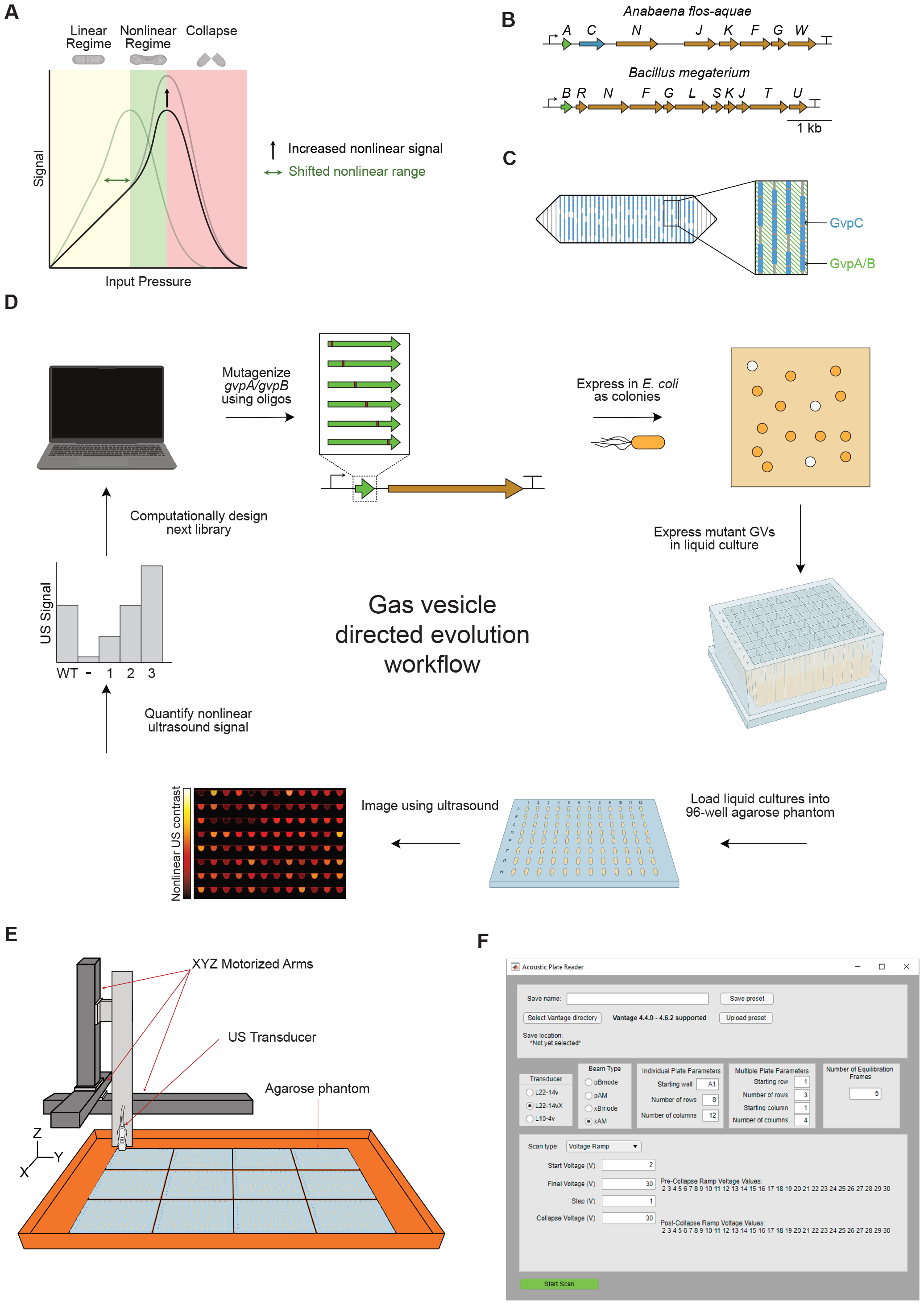
High-throughput directed evolution workflow for ARGs. (**A**) Three regimes of GV response to US. (**B**) Roles of the structural proteins GvpA/B and GvpC in GV structure. (**C**) Diagrams of the gene clusters used as starting points for evolution. (**D**) Schematic of directed evolution workflow for ARGs. The starting point GV structural protein is mutagenized, then expressed in *E. coli* as colonies on Petri dishes. Colonies that turn white are assumed to produce GVs, and are picked and expressed in liquid culture. Cultures of GV-expressing E. coli are then loaded into agarose phantoms and imaged using US at 15.625 MHz. The resulting nonlinear US intensity data are used to rank the performance of mutants and select the most promising ones for further mutagenesis. (**E**) Schematic of the Acoustic Plate Reader (APR), which is used for automated US image collection of up to 1152 samples of GV-expressing E. coli arrayed in 96-well agarose phantoms. (**F**) Image of the graphical user interface for the APR.

The primary GV structural protein – GvpA or its homolog GvpB – creates the cone-tipped cylindrical body of the GV, and optionally GvpC may attach to the outside of this structure and reinforce it mechanically (Figure 1B). It has already been shown that engineering GvpC to reduce its binding to GvpA can result in GVs with increased nonlinear signal or decreased collapse pressure,^19^ but GvpC serves as a limited target for engineering these phenotypes because not all GV types include GvpC. We chose to explore whether altering the primary structure of the main GV structural protein – GvpA in the *A. flos-aquae* cluster and GvpB in the *B. megaterium* cluster – could increase the amount of nonlinear US contrast produced by *E. coli* expressing either ARG type. We selected the GV gene clusters obtained from these species as our starting points based on the previous use of the *B. megaterium* cluster as a bacterial ARG^2^ and the use the *A. flos-aquae* cluster in reconstituted contrast agents and mammalian ARGs,^3,10,19^ making it desirable to obtain their efficient bacterial expression. Starting without the benefit of the recently published structures and structural models of these proteins,^20,21^ we chose an approach based on random mutagenesis and high-throughput acoustic screening of ARG mutants.

As starting points for evolution, we chose the minimal versions of the WT *B. megaterium* ATCC 19213 cluster^22^ (lacking *gvpA, gvpP*, and *gvpQ*) and the WT *A. flos-aquae* cluster (with only one copy of *gvpA*, and lacking *gvpV*) (Figure 1C). To engineer the desired nonlinear signal and collapse pressure phenotypes, we developed a method for high-throughput, semi-automated characterization of US contrast and GV collapse pressure in *E. coli* (Figure 1D).

First, we constructed scanning site saturation libraries of *gvpA* or *gvpB* in these clusters, and performed a selection for high levels of GV expression by inducing transformants on Petri dishes and picking only colonies that appeared white (GV-expressing bacteria appear white because GVs scatter light, in addition to US).^23,24^ These mutants were then expressed in liquid cultures in 96-well format and loaded into agarose phantoms. We imaged these phantoms using an automated scanning setup in which a software-controlled 3D translating stage raster-scans an US transducer above the submerged phantoms (Figure 1E), producing a set of US images in which samples with high GV expression appear bright. This pipeline allowed us to generate and acoustically screen several mutant libraries, from which we identified mutants with significantly enhanced acoustic phenotypes. We also created graphical user interfaces (GUIs) to simplify and standardize data acquisition (Figure 1F) and analysis. We termed this setup the “Acoustic Plate Reader” (Figure S1, Supplementary Video 1).

### Optimizing GV expression from WT A. flos-aquae and B. megaterium gene clusters

Before engineering the structural proteins, we optimized the expression of the WT *A. flos-aquae* and *B. megaterium* gene clusters in *E. coli* at 37°C. For each cluster, we cloned three origins of replication (ORIs) of different strengths (∼40, ∼20, and ∼5 copies/cell)^25^ (Figure 2A-B), and assessed their performance in liquid culture as a function of inducer concentration. For both clusters, the strongest ORI tested gave the highest nonlinear US signal (Figure 2C-D), and was chosen for future experiments. With the optimal ORIs selected for expression (Figure 2E-F), we then sought to optimize the autoinduction conditions to maximize nonlinear signal (in autoinduction media, increasing the concentration of glucose increases the cell density at which induction occurs, while increasing the concentration of the inducing sugar increases the level to which the transcription unit is induced). We performed titrations of glucose and arabinose and assessed the resulting nonlinear signal from the expressed constructs (Figure 2 G-H); we decided on concentrations of 0.25% glucose and 0.05% arabinose for induction of these constructs in future experiments, as these conditions yielded high GV expression from both constructs while leaving enough induction dynamic range to tune expression of mutants later without the need to alter any regulatory elements. We observed that US signal from the *A. flos-aquae* cluster peaked at a moderate arabinose concentration (Figure 2C and G), while expression from the *B. megaterium* cluster was highest at the maximum concentration (Figure 2D and H). We suspect that the signal decline from the *A. flos-aquae* cluster at high arabinose concentrations is due to the high metabolic burden associated with expressing so many non-native proteins in *E. coli*.

**Figure 2.**
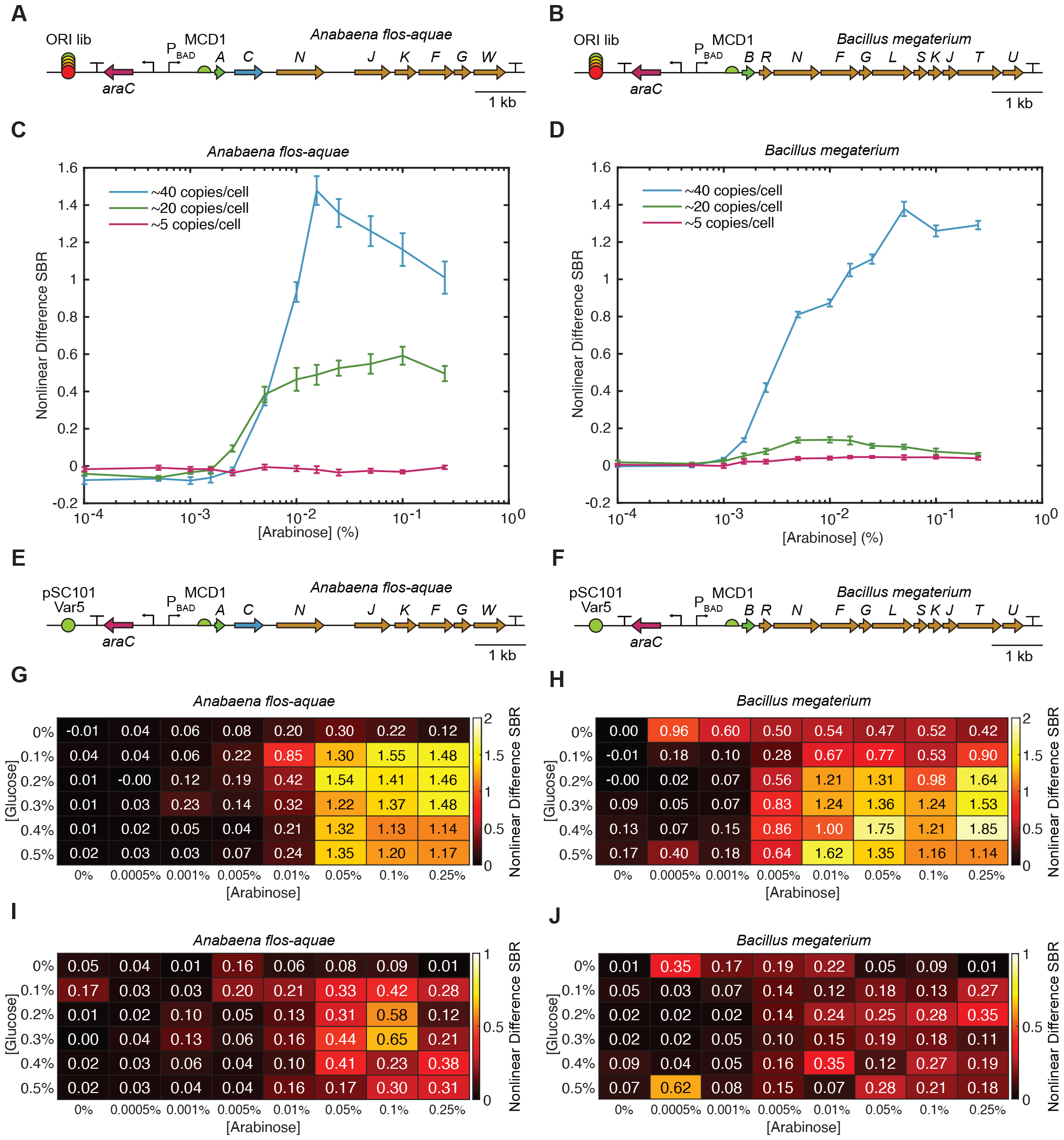
Optimization of GV expression from the WT *A. flos-aquae* and *B. megaterium* gene clusters. (**A-B**) Diagrams of the WT *A. flos-aquae* and *B. megaterium* gene clusters with libraries of origins of replication (ORIs) of different strengths. (**C-D**) Nonlinear US signal produced from expression of both clusters at three different copy numbers as a function of inducer concentration. The nonlinear difference SBR is the difference in signal-to-background ratio between pre- and post-collapse images of each sample (see Methods for details). Error bars represent standard error. N=8 biological samples (each an average of 3 technical replicates). (**E-F**) Diagrams of the optimized WT *A. flos-aquae* and *B. megaterium* gene clusters used for directed evolution, both of which used the pSC101-var5 ORI (∼40 copies/cell). (**G-H**) Mean and STD (I-J) nonlinear US signal produced by both WT clusters as a function of the concentrations of glucose and arabinose used for autoinduction. The concentrations selected for GV expression during library screening were 0.25% glucose and 0.05% arabinose. N=3 biological samples (each an average of 3 technical replicates).

### Round 1 directed evolution of A. flos-aquae GvpA and B.megaterium GvpB

To improve the nonlinear signal from the WT *A. flos-aquae* and *B. megaterium* clusters in *E. coli*, we designed scanning site-saturation libraries of the genes encoding the primary GV structural protein for each (*i.e*., *gvpA* for *A. flos-aquae*; *gvpB* for *B. megaterium*) (Figure 3A-B). This resulted in libraries containing 1400 and 1740 members for *gvpA* and *gvpB*, respectively (Table 1, Library Round 1). We constructed these libraries using a Golden Gate-based version of cassette mutagenesis,^26^ in which mutagenic oligonucleotides that tile the gene of interest are synthesized and cloned into an acceptor vector (Figure S2A-B; see methods for details). We chose this approach over error-prone PCR because of its ability to generate defined libraries which have a controllable number of mutations per member and which lack unwanted mutants (*i.e*., premature stop codons and multiple codons that code for the same mutant).

**Figure 3.**
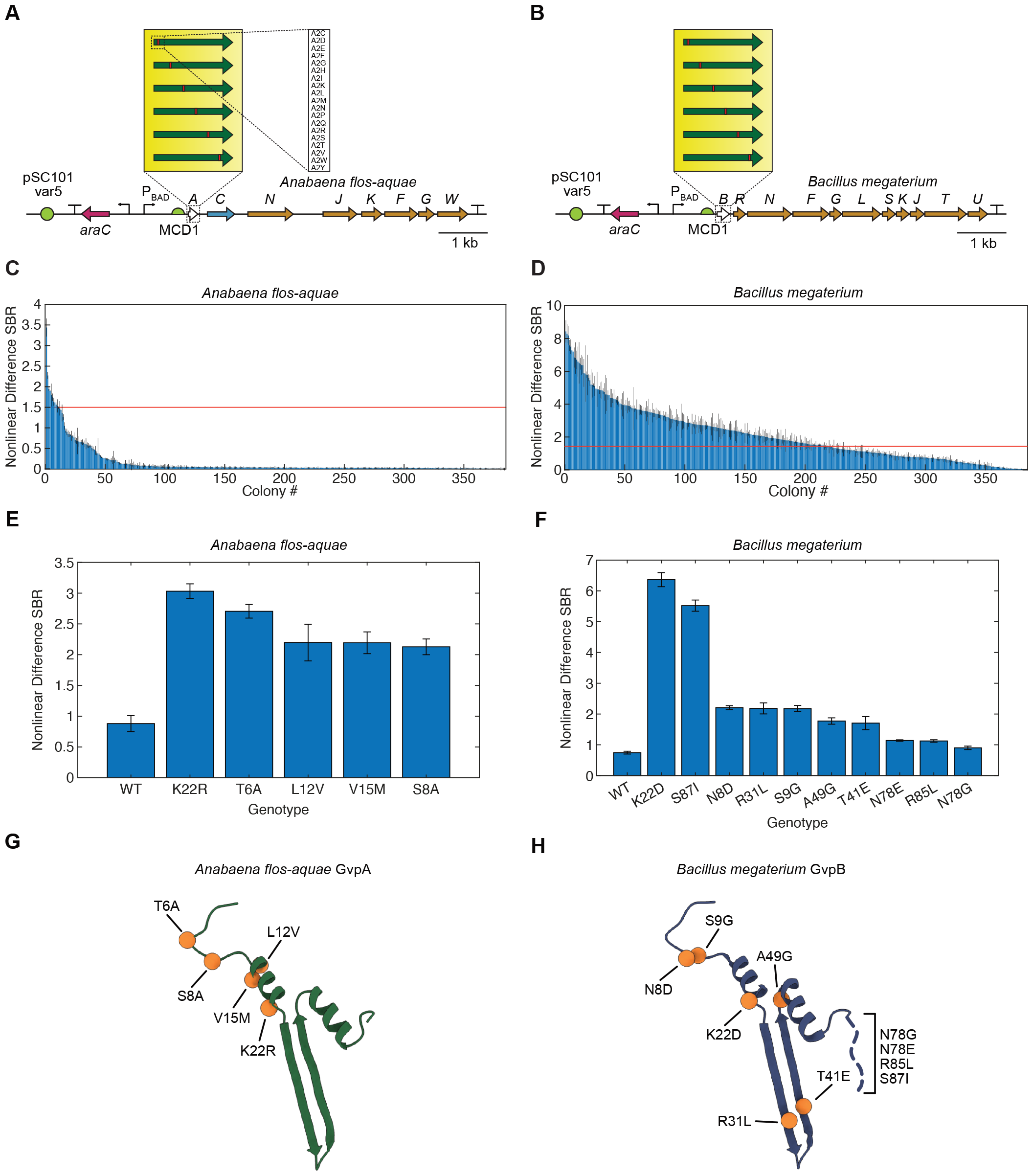
First round of directed evolution of *A. flos-aquae* and *B. megaterium* structural proteins. (**A-B**) Diagrams of the mutagenized *A. flos-aquae* and *B. megaterium* gene clusters, depicting the scanning site saturation libraries screened in the first round of evolution. (**C-D**) Nonlinear US difference signal-to-background ratio (SBR) from all screened mutants of both clusters. Red lines indicate the difference SBR of the WT for that cluster. Error bars represent standard error. N=3 technical replicates of one biological sample. (**E-F**) Nonlinear US difference SBR for the WT and top mutants for each cluster. Error bars represent standard error. N=4 biological samples (each an average of 3 technical replicates). (**G-H**) Locations of top mutations from E-F in the GvpA/GvpB structure (PDB 8GBS and 7R1C).

When induced in solid culture, these libraries produce three distinct types of colonies: 1) blue colonies, in which the dropout chromoprotein was not excised during assembly, returning the original acceptor vector; 2) low-opacity colonies that lack GV expression or express small amounts, either because they contain a mutant that reduces GV expression or because the mutant gene did not insert correctly during assembly; 3) high-opacity colonies with high GV expression. Colony opacity corresponds to GV expression because the low index of refraction of air inside GVs relative to surrounding aqueous media results in light scattering.^23,27^ We used this readout to select only the mutants with high GV expression for further study. We then expressed these mutants (384 from each of the two libraries) in 96-well liquid cultures, and imaged them in the APR in 96-well agarose phantoms (Figure 1D and Figure S1). Among the GvpA mutants, only a small number showed significantly higher nonlinear US signal than the WT (Figure 3C), while many GvpB mutants showed an increase (Figure 3D). This was likely because the GvpA construct fails to produce strongly opaque colonies when grown in solid culture, making it impossible to enrich for functional mutants prior to US screening; thus, the mutants screened via US from the GvpA library represent a random subset of the library, while those from the GvpB library are enriched for GV-producing sequences.

We chose up to 10 unique mutants with the highest US signal from each library and re-cloned them (see methods) for validation and further characterization of their nonlinear acoustic signal (Figure 3E-F and Figure S3A-B) and OD600 (Figure S3C-D). The top two mutants from each library — GvpA-T6A and -K22R, and GvpB-K22D and -S87I — generated nonlinear US signals 3.07-, 3.44-, 8.54-, and 7.41-fold higher than their parents, respectively, while growing to similar densities in liquid culture. The mutations found in the top 5 and top 10 variants from the GvpA and GvpB libraries, respectively, are shown in Figure 3G-H. These mutations cluster in the N-terminal linker and bridge domains, as well as the hinge and wall domains, and the C-terminal tail.^20,21^ Notably, no mutations occur in the C-terminal stabilization domain.

### Round 2 directed evolution of A. flos-aquae GvpA and B. megaterium GvpB

We next performed a second round of directed evolution on these clusters by generating three distinct libraries: two scanning site saturation libraries of the top two mutants of *A. flos-aquae gvpA* (T6A and K22R) and a paired recombination library of the top 10 unique mutants of *B. megaterium gvpB* (Figure 4A-B) (though some members of this library contained three mutations due to a well-documented issue with amplifying oligonucleotide pools; see Methods for explanation). We cloned and screened these libraries using the same methods described for the first round of evolution (Figure 1D), and identified several mutants with greatly improved signal over their parents in both libraries (Figure 4C-D).

**Figure 4.**
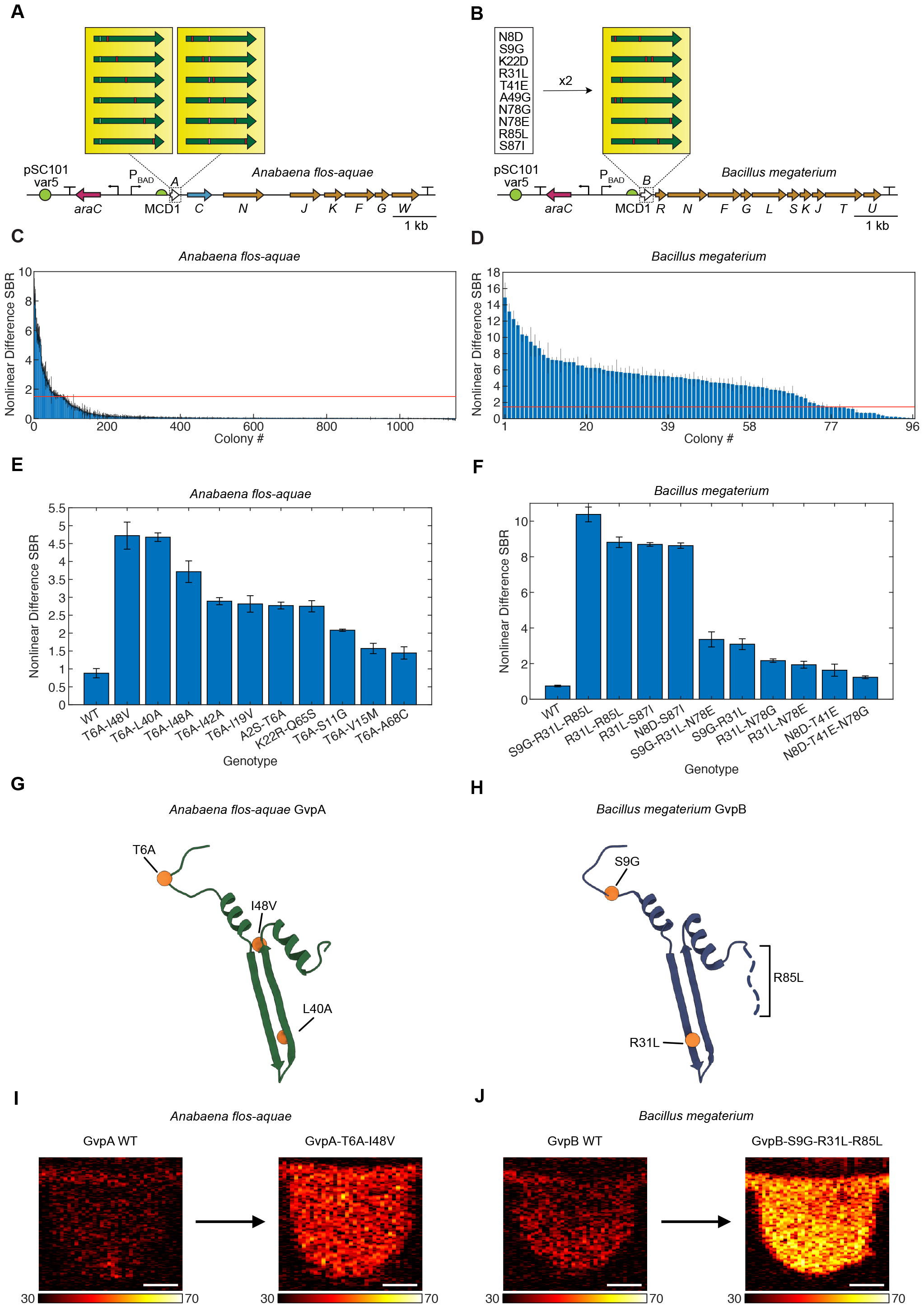
Second round of directed evolution of *A. flos-aquae* and *B. megaterium* structural proteins. (**A-B**) Diagrams of the mutagenized *A. flos-aquae* and *B. megaterium* gene clusters used in the second round of evolution. The best two mutants of *A. flos-aquae* gvpA were used as parents for another scanning site saturation library, and the best ten mutants of *B. megaterium* gvpB (listed in figure) were used to create a paired recombination library. (**C-D**) Nonlinear US difference signal-to-background ratio (SBR) from all screened mutants of both clusters. Red lines indicate the difference SBR of the WT for that cluster. Error bars represent standard error. N=3 technical replicates of one biological sample. (**E-F**) Nonlinear US difference SBR for the WT and top ten mutants for each cluster. Error bars represent standard error. N=4 biological samples (each an average of 3 technical replicates). (**G-H**) Locations of mutations from the top mutants from E-F in the GvpA/GvpB structure. (PDB 8GBS and 7R1C) (**I-J**) Representative nonlinear US images of the brightest mutants identified in this study, as well as their respective WT parents. Scale bars 1 mm.

We characterized the top 10 unique mutants from each library in terms of their nonlinear acoustic signal (Figure 4E-F and Figure S4A-B) and OD600 (Figure S4C-D), and identified GvpA-T6A-L40A, GvpA-T6A-I48V, and GvpB-S9G-R31L-R85L as the top-performing variants. These mutants generated nonlinear signals 5.32-, 5.37-, and 13.93-fold higher than their parents, respectively, while allowing the bacteria expressing them to grow to similar densities in liquid culture. The mutations found in the top 2 and top 1 variants from the second-round GvpA and GvpB libraries, respectively, are shown in Figure 4G-H. Similar to the mutations identified from the first-round libraries, these mutations cluster in the N-terminal linker domain, as well as the hinge and wall domains, and the C-terminal tail, but not the C-terminal stabilization domain.^20,21^ Representative nonlinear US images of GvpA-T6A-L40A and GvpB-S9G-R31L-R85L, as well as the WT parents, are shown in Figure 4I-J. In addition to showing increased nonlinear contrast (Figure S5A-B), the top variants have slightly higher collapse pressure than their WT parents when normalized for nonlinear contrast (Figure S5C-D).

### Expression characteristics of top mutants

We performed whole-cell transmission electron microscopy (TEM) on *E. coli* expressing either WT or mutant ARGs to evaluate changes in expression levels. TEM revealed that these mutations increased the expression levels of both ARG types, either by increasing both the typical and maximum number of GVs per cell (in the case of GvpA-T6A-L40A and GvpA-T6A-I48V) or by making the number of GVs expressed per cell more consistent across all cells in the culture (in the case of GvpB-S9G-R31L-R85L) (Figure 5A-E, Figure S6-7).

**Figure 5.**
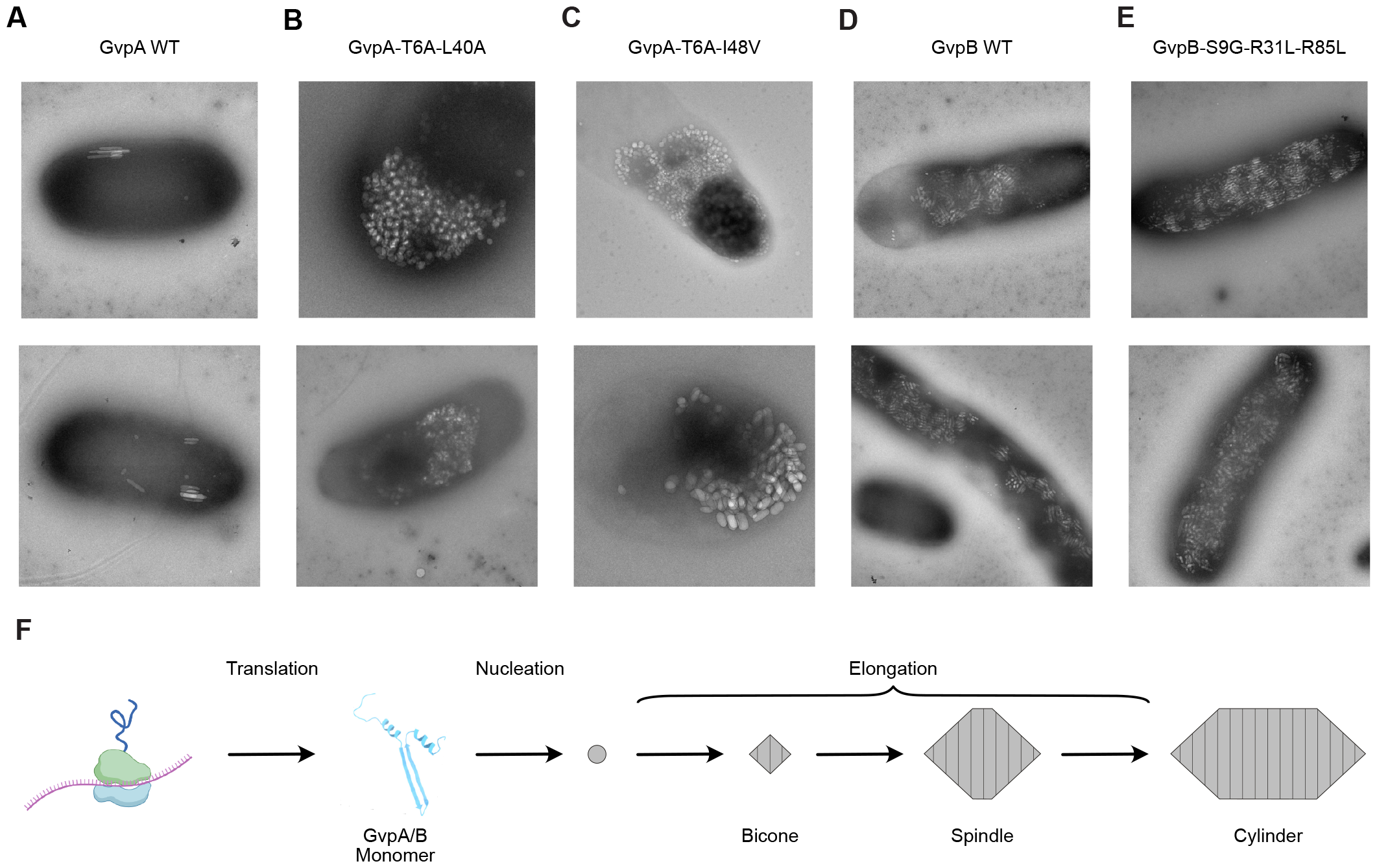
TEM of *E. coli* after expressing top-performing *A. flos-aquae* GvpA and *B. megaterium* GvpB mutants. (**A-E**) TEM images of WT and mutant GVs expressed in *E. coli*. (**F**) Diagram of the GV formation process.

## DISCUSSION

Our results establish the first method for high-throughput, semi-automated acoustic screening of biomolecules expressed in cells. When used to evolve two ARG clusters—those from *A. flos-aquae* and *B. megaterium*—this method yielded ARG constructs which show 5- to 14-fold improvements in their nonlinear acoustic scattering.

The mutations identified in this study appear to increase nonlinear US signal by increasing the maximum number of GVs produced per cell and/or making GV production more consistent across a cell population. These changes could be due to improved expression of GvpA/GvpB monomers or their incorporation into growing GVs (Figure 5F). In addition, it is possible that some mutations contribute to increased nonlinear scattering of individual GVs by altering their mechanical properties.^14,17,28,29^

These results represent a major advance in the way that acoustic biomolecules can be engineered. In the same way that high-throughput screening tools such as plate readers and flow cytometry enabled the engineering of fluorescent proteins and the many sensors derived from them by dramatically increasing the sizes of libraries that can be screened in these experiments, so too will the increased throughput, reliability, and standardization introduced by the Acoustic Plate Reader enable the engineering of next-generation ARGs and their derivatives.

While these evolved constructs represent substantial improvements over their parents, further improvements are required. First, both ARGs could benefit from further improvements in nonlinear contrast; this will likely be achieved through a combination of protein engineering (including not only the structural proteins engineered in this study, but also the assembly factors that assist in GV formation) and expression tuning (ORI, RBSs, and promoter) aimed at increasing both the amount of nonlinear contrast produced per GV and the number of GVs produced per cell. Relatedly, it would be desirable to engineer GVs with higher collapse pressures or ones whose collapse pressure is unchanged while having a significantly lower buckling threshold.

Additional engineering is needed to ensure the mutational stability of these constructs for *in vivo* applications, for example through chromosomal integration or inclusion of plasmid stability elements.^4^ APR screening could facilitate any tuning required at the transcriptional (promoter) and translational (RBS) levels. Such tuning would potentially make the more compact *A. flos-aquae* and *B. megaterium*-derived ARGs competitive with the larger *Serratia*-derived ARGs, which currently provide the best *in vivo* imaging performance.^4^ To further accelerate ARG development, we need a deeper understanding of how mutations to GvpA/GvpB affect both their structures^20,21^ and the protein-protein interactions in which they participate during GV assembly^30–33^, as well as biochemical methods to characterize intermediate steps that cannot be assayed by ultrasound, such as GV nucleation (Figure 5F).

By enabling the large-scale generation and high-throughput acoustic screening of ARG variants, the APR and its associated protocols allow the suite of modern protein engineering techniques to be applied to ARGs.

## MATERIALS AND METHODS

### Plasmid construction via MoClo

The EcoFlex MoClo system^37^ was used to create all vectors cloned in this study, including existing parts (Addgene Kit # 1000000080) and custom-made parts (Table 2). Custom-made parts were introduced into the existing EcoFlex system as follows: 1) ORIs were selected from the pSC101-varX series^25^; promoters were selected from the Marionette series^38^; RBSs were selected from the MCDX series^39^; terminators were selected from the ECK and LXSXPX series^40^; 2) parts were either synthesized as fragments (Twist Bioscience) and subsequently PCRed using Q5 (NEB), or synthesized as duplex oligos (IDT); 3) parts were cloned into the corresponding part acceptor vector (Table 2) via Golden Gate to ensure that they received the appropriate assembly overhangs. EcoFlex assemblies were conducted as described in Supplementary Note 1 and electroporated into NEB Stable E. coli (except for the MetClo-based library acceptor vectors, which were transformed into DH10B-M.Osp807II^41^). Transformations were recovered for 2 hr in 1 mL of SOC at 37°C and 250 RPM, and plated on Petri dishes containing Lennox LB with 1% agar, 100 ug/mL kanamycin, and 1% glucose (for catabolite repression of the PBAD promoter). Colonies were picked into 1.5 mL liquid cultures of Lennox LB with 100 ug/mL kanamycin and 1% glucose in 96-well format and grown overnight to saturation. These cultures were then miniprepped using reagents and a protocol from Qiagen, a lysate clearing plate from Bio Basic (SD5006), and a DNA-binding plate from Epoch Life Sciences (2020-001). All constructs were verified by whole-plasmid sequencing (Primordium Labs).

### Liquid culture GV expression in E. coli

GVs were expressed in E. coli liquid cultures in 96-well format according to the following general protocol, with modifications for specific experiments described below.

Miniprepped DNA was electroporated into NEB Stable *E. coli*, and transformations were recovered for 2 hr at 37°C in 1 mL of SOC. Transformations were then inoculated at a dilution of 1:100 into autoinduction Lennox LB containing 100 µg/mL kanamycin, 0.6% glycerol, and the appropriate concentrations of glucose and inducer for the experiment (see below). These expression cultures were set up in 500 uL volumes in deep-well 96-well plates (square wells used for maximum culture aeration; USA Scientific 1896-2800) sealed with porous tape (Qiagen 19571) and incubated at 37°C and 350 RPM for 20 hr. Cultures were stored at 4C until being loaded into phantoms for Acoustic Plate Reader scans. For the concentrations of glucose and arabinose described below, in experiments where titrations were used, 100X stocks of these sugars were prepared in 1X PBS and diluted 1:100 into the cultures when setting up the experiments.

The following concentrations were used for the ORI titration experiments shown in Figure 2A-D: glucose: 0.25%; arabinose: 0, 0.0005, 0.001, 0.00155, 0.0025, 0.005, 0.01, 0.0155, 0.025, 0.05, 0.1, 0.25%.

The following concentrations were used for the parent expression optimization experiments shown in Figure 2E-H: glucose: 0, 0.1, 0.2, 0.3, 0.4, 0.5%; arabinose: 0, 0.0005, 0.001, 0.005, 0.01, 0.05, 0.1, 0.25%.

The following modifications were made for the library screening experiments shown in Figure 3A-C and 4A-C: 1) assembled libraries were transformed multiple times into NEB Stable *E. coli*, and it was ensured that the number of transformants produced was at least 100X the number of unique sequences expected in the library; 2) prior to expression in liquid culture, libraries were expressed in solid culture as colonies on Lennox LB with 100 ug/mL kanamycin, 0.6% glycerol, 0.25% glucose, and 0.05% arabinose at a density of ∼100 colonies/dish. Colonies were grown for 48 hr at 37°C, and 380 opaque colonies were picked for each library, as well as 4 colonies for the library’s parent, into the wells of 96-well PCR plates containing 100 uL of Lennox LB with 100 µg/mL kanamycin and 1% glucose, and grown to saturation overnight at 30°C. These saturated liquid cultures, rather than transformations, were used to set up expression cultures as described above; 3) 0.25% glucose, and 0.05% arabinose were used to induce expression in these experiments.

The following concentrations were used for the mutant expression experiments shown in Figure 3E-H and 4E-H: glucose: 0.25%; arabinose: 0.05%.

The following concentrations were used for the multiplexing experiments shown in Figure 5A-B: glucose: 0.25%; arabinose: 0.05%.

### Scanning site saturation and recombination library generation

Scanning site saturation libraries were generated via a Golden Gate-based version of the cassette mutagenesis strategy previously described.^42^ Briefly, the *A. flos-aquae* GvpA and *B. megaterium* GvpB coding sequences were divided into sections that tiled the gene, and oligos were designed to have a variable middle region with flanking constant regions against which PCR primers were designed (these primers also contain the evSeq^43^ inner adapters for optional deep sequencing of the library). Depending on the library being created (*i.e*., scanning site saturation or recombination), the variable region was designed to either sequentially saturate each residue or recombine pairs of the mutations listed in Figure 4B (mutations identified during screening of the first round of scanning site saturation GvpB). The MATLAB scripts used to generate the oligo sequences for both the scanning site saturation and recombination libraries are available in Supplementary Code 1, and the oligo sequences themselves are listed in Table 1. Oligos were synthesized as a pool by Twist Biosciences or Integrated DNA Technologies, and were amplified by PCR (both to make them double-stranded and to generate enough DNA for Golden Gate assembly) using KAPA HiFi HotStart ReadyMix according to the manufacturer’s instructions, but with 10 cycles, 100 ng of oligo pool template, and 1 uM of each primer. PCR products were run on a 2% agarose gel and purified using Qiagen reagents according to the manufacturer’s instructions, but with a 5 uL final elution volume of water. Fragments were then assembled with the corresponding library acceptor vector (Table 2) in a Golden Gate reaction using reagents from New England Biolabs according to the manufacturer’s instructions. Assemblies were then expressed (first in solid culture and then in liquid culture) according to the protocol above.

It is important to note that oligo pools whose members have very high sequence similarity (as was the case in the pools used in this study, in which members differed by only a few bp) have a high likelihood of mutation swapping during PCR which increases with the number of cycles used. The manufacturer proposes that this is due to template swapping from one cycle to the next between incompletely-copied strands. We notice this often in our libraries (*i.e*., libraries synthesized to have two mutations per member would contain a small number of sequences with zero or three mutations per member after PCR), and we minimized the number of PCR cycles used to amplify these libraries. However, some of the best round 2 GvpB mutants contained three mutations for this reason.

### Acoustic Plate Reader scans

The general protocol for preparing and scanning liquid cultures samples of GV-expressing E. coli in 96-well format is described in Figure S1 and the corresponding figure caption. Detailed instructions on how to build and use this system, as well as troubleshooting and bug-reporting information, are provided at https://github.com/shapiro-lab/acoustic-plate-reader.

The specific US pulse sequence parameters used for collecting the data shown in each figure are presented in Table 3.

For pre-/post-collapse and voltage ramp scans, the nonlinear difference SBR was calculated as: [(pre-collapse sample mean) - (post-collapse sample mean)] / (post-collapse background mean), where means are calculated from the nonlinear signal in a region of interest containing either the sample or an empty region of the phantom. For voltage ramp scans, this quantity was calculated for each pre-collapse image; for simple pre-/post-collapse scans, this quantity was calculated only once for the single pre-collapse image. Importantly, in all cases the two images being compared in each calculation were acquired at the same voltage (*i.e*., the pre- and post-collapse images were collected under the same imaging conditions).

For collapse ramp scans, the nonlinear SBR was calculated as: (sample mean) / (background mean), where means are calculated from the nonlinear signal in a region of interest containing either the sample or an empty region of the phantom. This quantity was calculated for each image at each voltage.

### Validation of best mutants

Selected mutants from each library were miniprepped and sequenced as described above. Unique mutants were then re-cloned using MoClo (see above) before undergoing validation testing to avoid the possibility that these plasmids accrued expression-reducing mutations during the GV expression steps performed during library screening. To prepare fragments for these MoClo assemblies, *gvpA*/*gvpB* mutant CDSs were PCRed using the primers described in Table 4 (which were selected based on the sequence of the mutant being amplified) and prepared for Golden Gate assembly as described above.

### OD600 measurements

OD 600 culture measurements were performed on a Tecan Spark plate reader using the “Absorbance” protocol with the following settings: 600 nm measurement wavelength, 10 flashes, 50 ms settle time. Measurements were collected for 200 uL of culture and normalized to a 1 cm path length using the built-in “Pathlength Correction” feature.

### Negative stain TEM imaging

Three microliters of *E. coli* culture expressing GVs were applied to a freshly glow-discharged (Pelco EasiGlow, 15 mA, 1 min) Formvar/carbon-coated, 200-mesh copper grid (Ted Pella), and then incubated for 1 minute. Excess solution was blotted with filter paper, and the grids were washed three times with buffer (20 mM HEPES buffer; pH 7.5, 100 mM NaCl). Subsequently, the sample was stained with a 2% uranyl acetate solution for 1 min, blotted, and air-dried. Images were acquired using a Tecnai T12 electron microscope (FEI, now Thermo Fisher Scientific) operating at 120 kV and equipped with a Gatan Ultrascan 2k × 2k CCD.

## DATA AVAILABILITY

Selected plasmids are available through Addgene (202023, 202024, 202025). Detailed instructions on how to build and use the Acoustic Plate Reader, as well as troubleshooting and bug-reporting information, are provided at https://github.com/shapiro-lab/acoustic-plate-reader. All other data and code are available from the corresponding author upon reasonable request.

## ACKNOWLEDGEMENTS

The authors would like to thank Rohit Nayak for providing calibration data for the US transducer used for imaging. Transmission electron microscopy was done in the Beckman Institute Resource Center for Transmission Electron Microscopy at Caltech. This research was supported by the National Institutes of Health (R01-EB018975 to M.G.S.), the Chan-Zuckerberg Initiative and Pew Charitable Trust. R.C.H. was supported by the Caltech Center for Environmental Microbial Interactions. Related research in the Shapiro Laboratory is supported by the David and Lucile Packard Foundation. M.G.S. is an investigator of the Howard Hughes Medical Institute.

## AUTHOR CONTRIBUTIONS

R.C.H., Z.J., and M.G.S. conceived and designed the study. Z.J. and D.R.M. designed and built the Acoustic Plate Reader hardware. Z.J., D.P.S., and D.R.M. wrote the MATLAB scripts for data acquisition with the Acoustic Plate Reader. R.C.H., Z.J., and D.P.S. wrote the MATLAB scripts for data analysis from the Acoustic Plate Reader. M.S. designed the MATLAB graphical user interfaces for Acoustic Plate Reader data acquisition and analysis. R.C.H., M.S., K.W., H.K.S., R.D., and R.Z. performed directed evolution experiments. P.D. performed TEM imaging. R.C.H. analyzed all data. R.C.H. wrote the paper, with input from all authors. M.G.S. supervised the research.

## COMPETING INTERESTS

The authors declare no competing financial interests.

## SUPPLEMENTARY INFORMATION

Supplementary Figures 1-7

Supplementary Note 1

Supplementary Video 1

Supplementary Tables 1-4

## SUPPLEMENTARY INFORMATION

**Figure S1:**
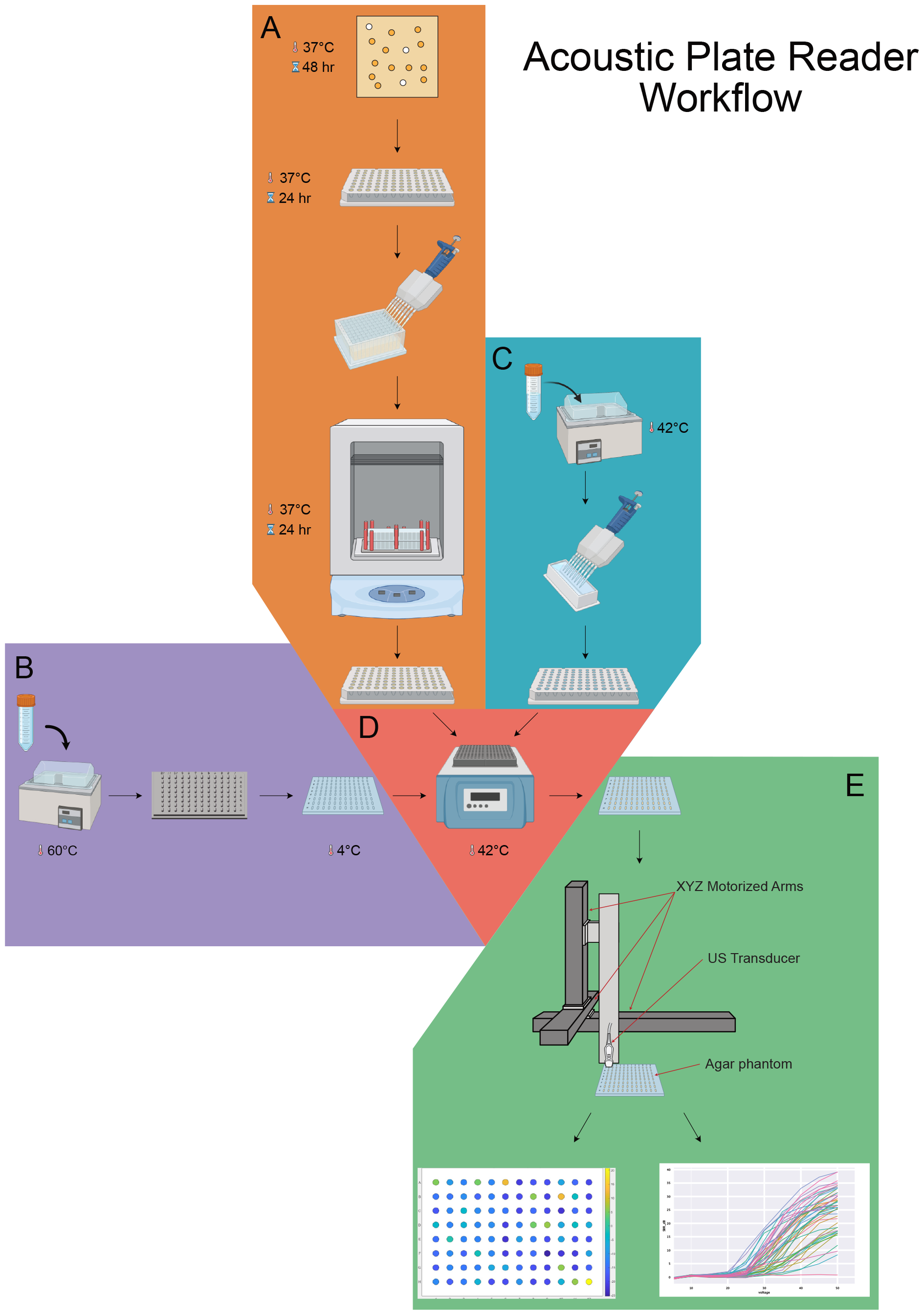
Detailed diagram of the acoustic plate reader workflow. (**A**) GVs are expressed in *E. coli* as colonies on Petri dishes for 48 hr at 37°C, then colonies are picked into LB and grown to saturation in liquid culture for 24 hr at 37°C. These saturated liquid cultures are then diluted 1:100 into autoinduction LB and expressed for 24 hr at 37°C in 500 uL cultures in deep-well 96-well plates (square wells used for maximum culture aeration; USA Scientific 1896-2800). Aliquots of these cultures are aliquoted into an un-skirted 96-well PCR plate for subsequent loading into phantoms. (**B**) A solution of 2% Ultrapure Agarose (Invitrogen, 16500500) is prepared in 1X PBS and incubated at 60C for at least 12 hr to degas. Agarose phantoms are then made by pouring 75 mL of this solution into a 96-well phantom mold and incubating at 4C for 10 min. (**C**) A solution of 1% low-melting-temperature agarose (Goldbio, A-204-100) is prepared in 1X PBS and incubated at 60C for at least 12 hr to degas. This solution is then aliquoted into an un-skirted 96-well PCR plate to be used for phantom loading. (**D**) Phantoms from B are loaded by placing the 96-well PCR plates from A and C into 96-well heat block at 42C, and combining equal volumes of culture and agarose before pipetting into the empty phantom. (**E**) Phantoms from D are scanned using the acoustic plate reader, which generates US data for each sample and can image up to 12 96-well phantoms in a single scan.

**Figure S2:**
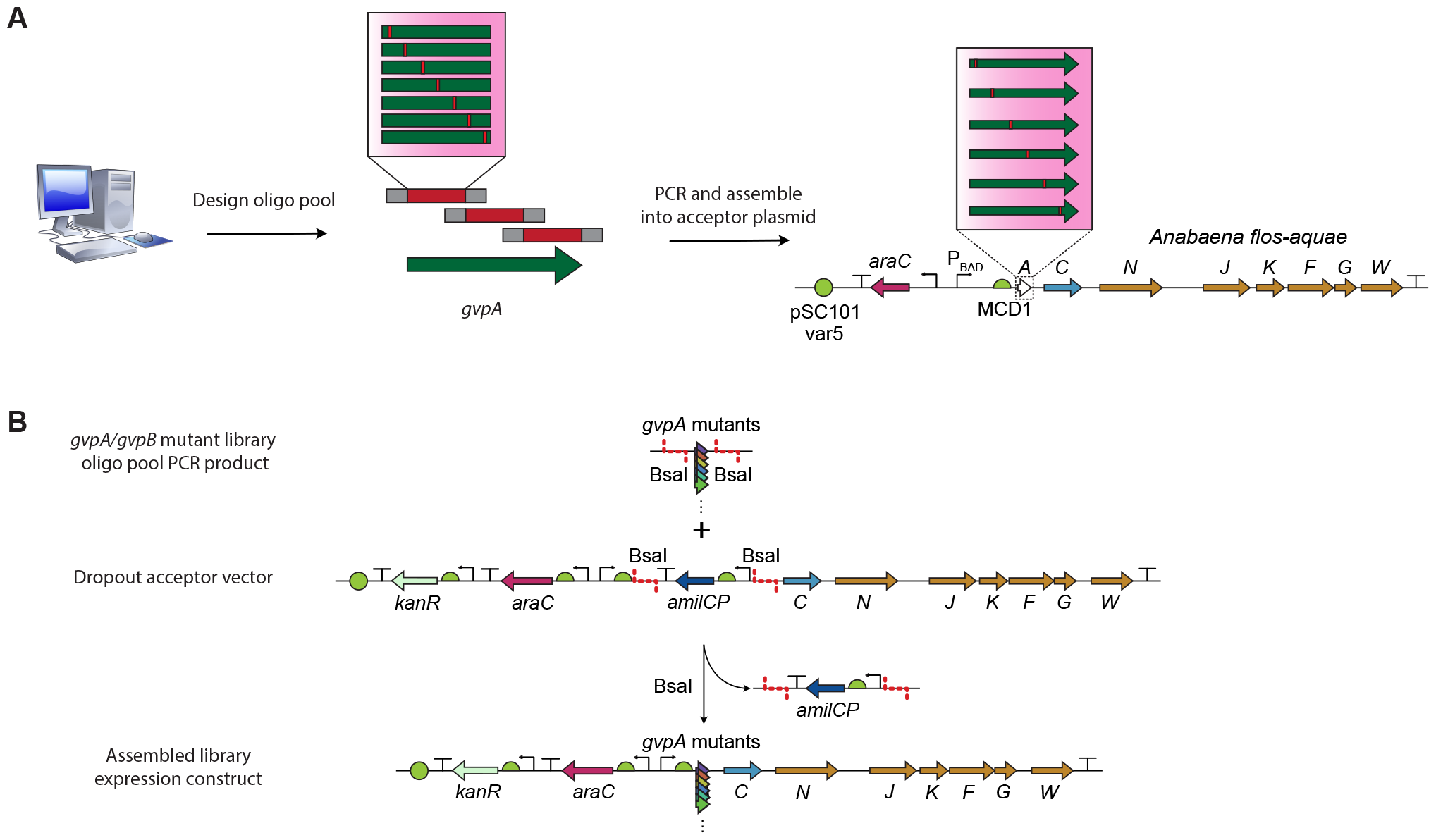
Details of *gvpA*/*gvpB* mutant library construction and screening. (**A**) Overview of workflow for creating either scanning site saturation or recombination libraries. (**B**) Details of library assembly via a Golden Gate-based version of cassette mutagenesis (see Methods).

**Figure S3:**
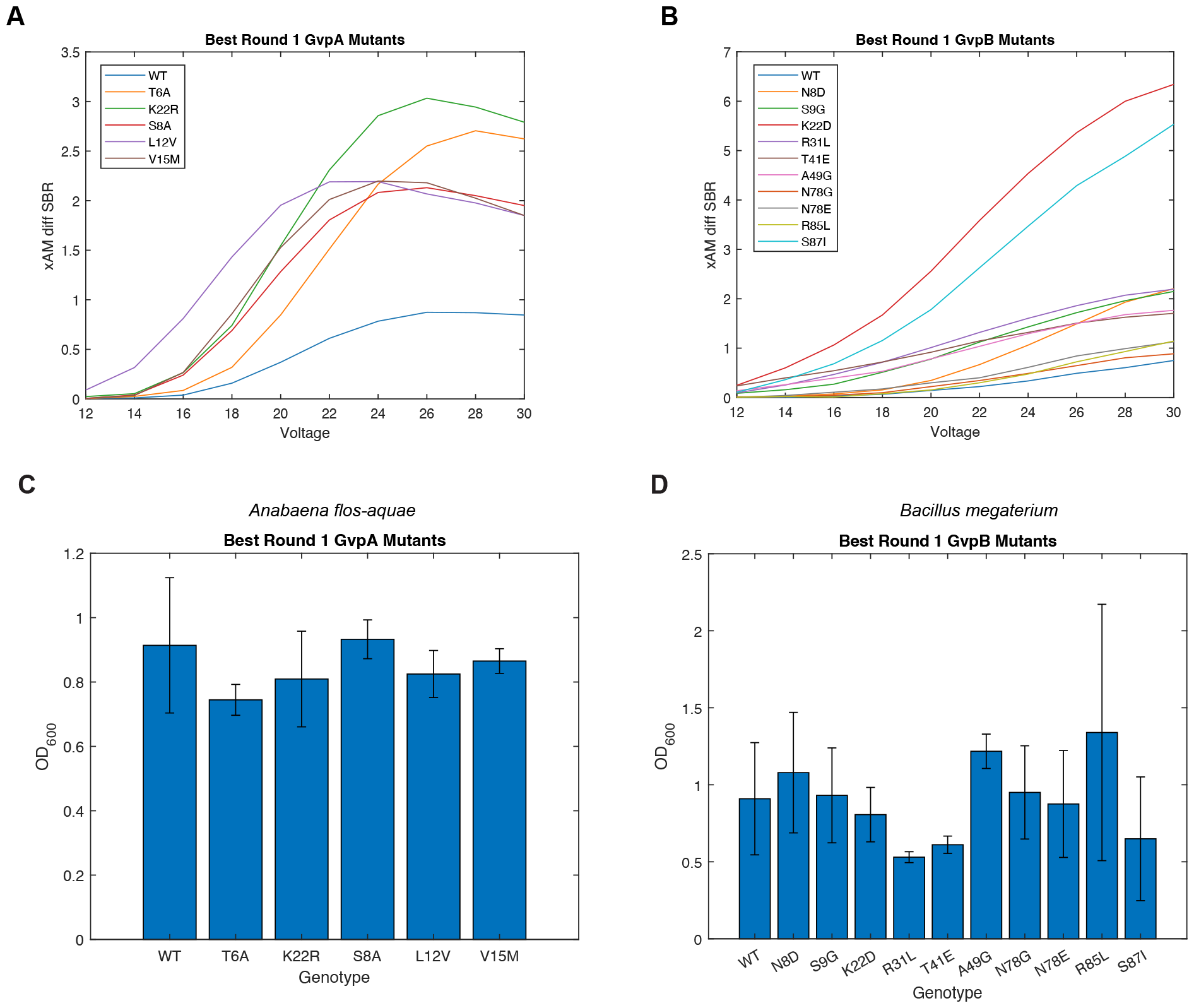
Characterization of the top mutants from Round 1 of evolution. (**A-B**) xAM difference SBR as a function of pressure for each of the top mutants. N=4 biological samples (each an average of 3 technical replicates). (**C-D**) OD600 measurements for the mutants shown in A-B. N=4 biological samples.

**Figure S4:**
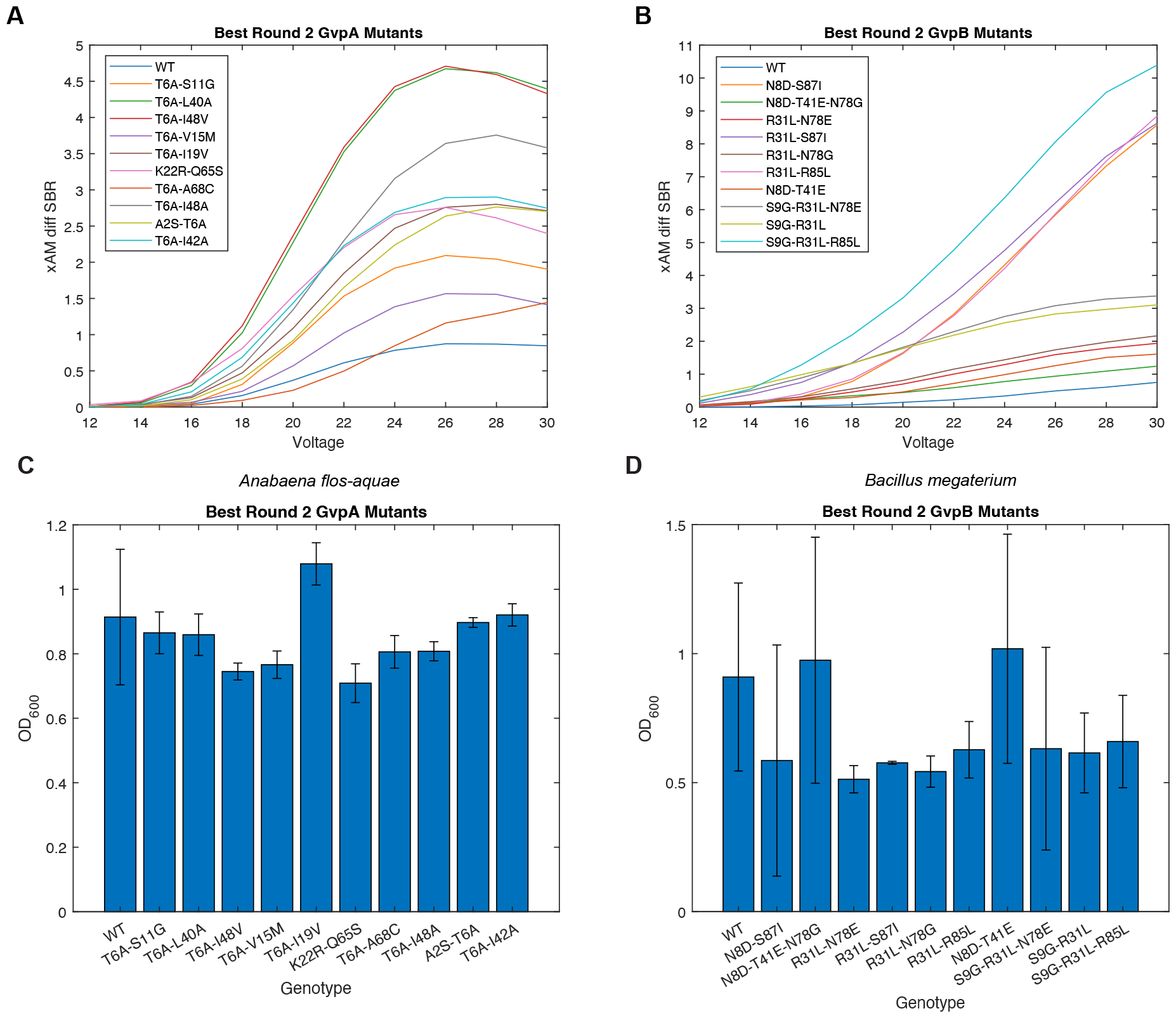
Characterization of the top mutants from Round 2 of evolution. (**A-B**) xAM difference SBR as a function of pressure for each of the top mutants. N=4 biological samples (each an average of 3 technical replicates). (**C-D**) OD600 measurements for the mutants shown in A-B. N=4 biological samples.

**Figure S5:**
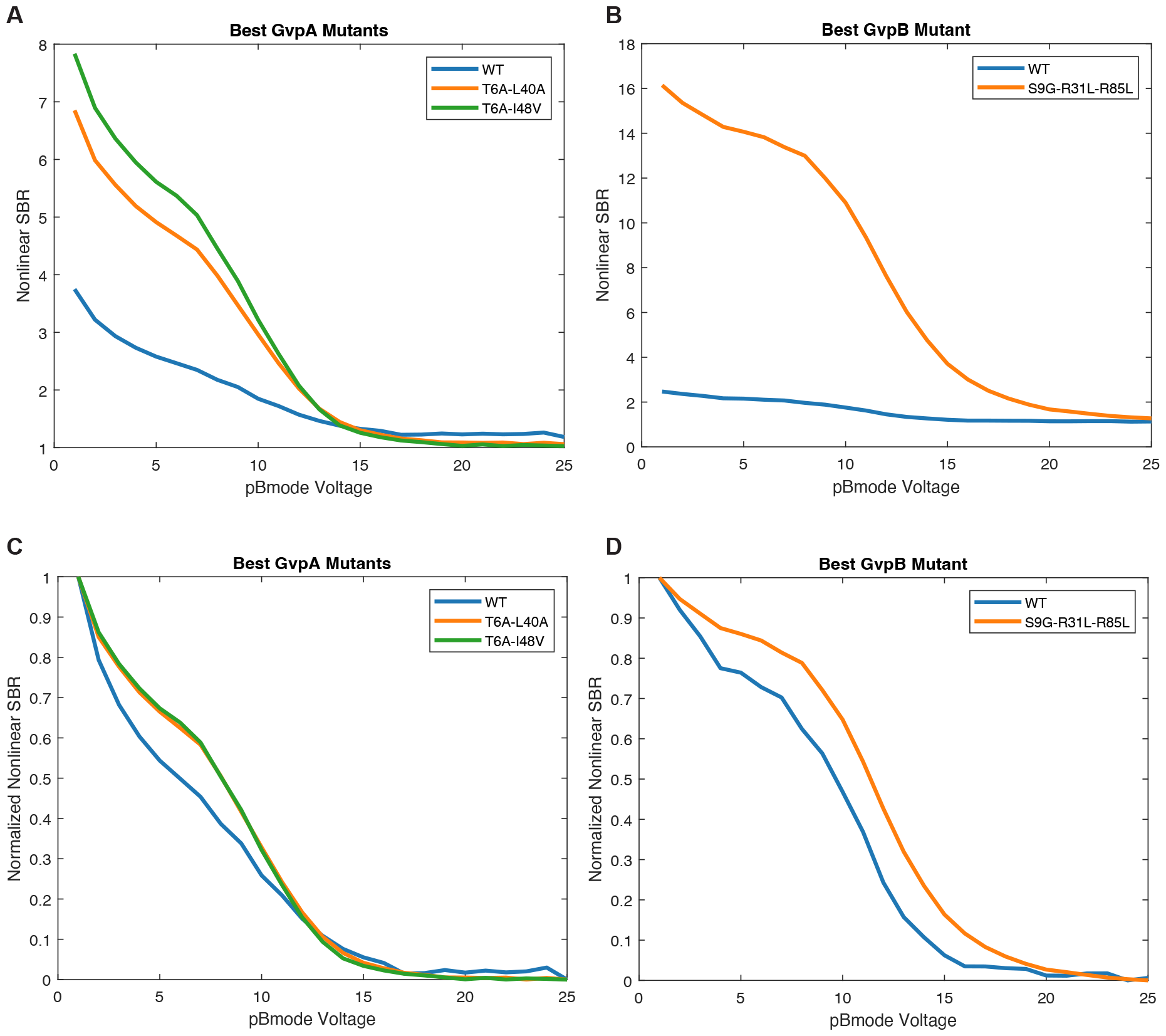
Acoustic collapse pressure curves for the best mutants identified in this study. (**A-B**) xAM acoustic collapse pressure curves for the top-performing mutants identified in this study. (**C-D**) Data from A-B normalized to the same min and max. N=4 biological samples.

**Figure S6:**
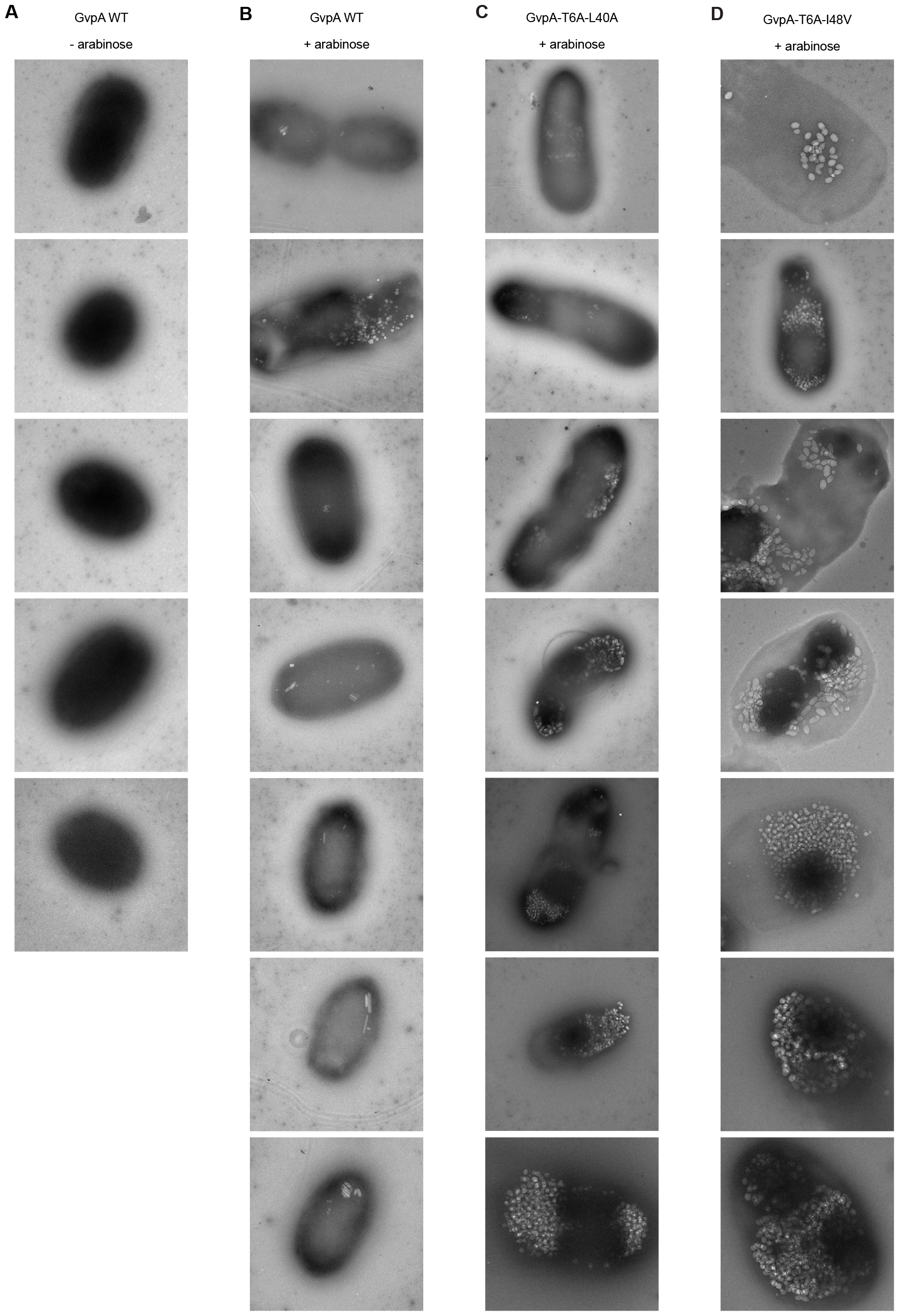
TEM images of *E. coli* cells expressing WT or mutant *A. flos-aquae* GVs. For each sample, 5-25 images were collected; a representative set is shown, ordered from least to most GVs produced per cell.

**Figure S7:**
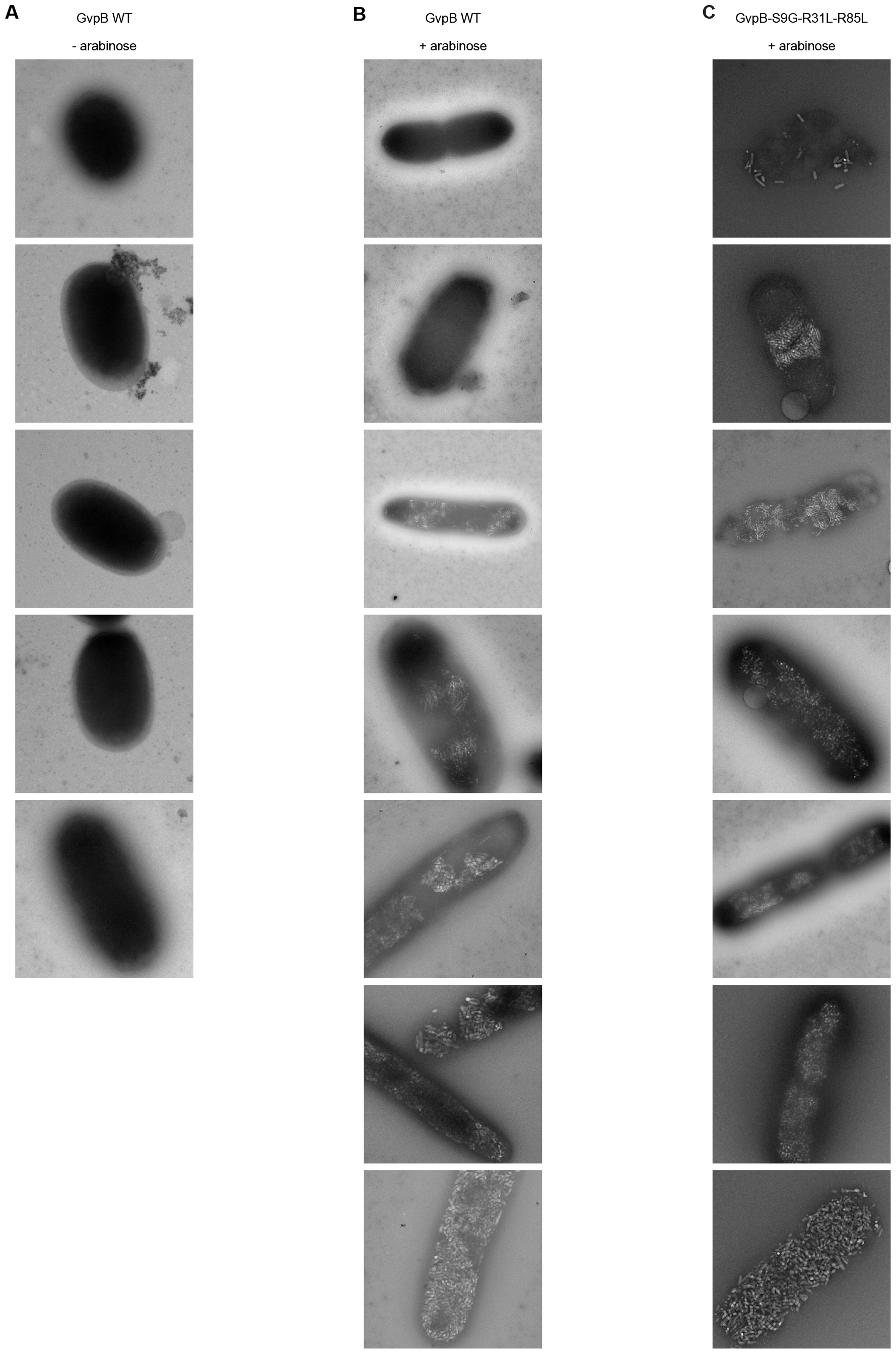
TEM images of *E. coli* cells expressing WT or mutant *B. megaterium* GVs. For each sample, 5-25 images were collected; a representative set is shown, ordered from least to most GVs produced per cell.

## Supplementary Note 1: Golden Gate reactions

### Master mix recipes

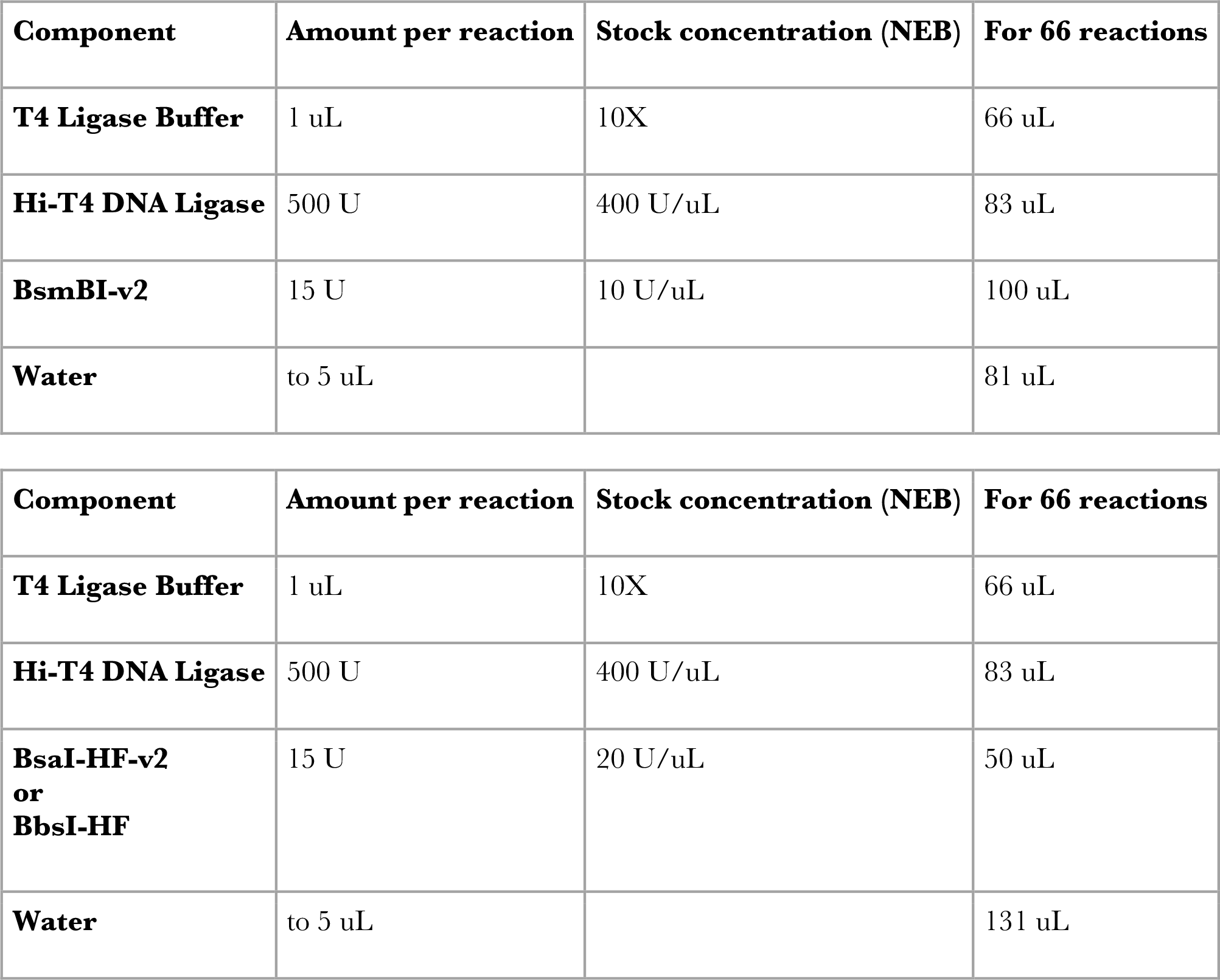

| Component | Amount per reaction | Stock concentration (NEB) | For 66 reactions |
| --- | --- | --- | --- |
| **T4 Ligase Buffer** | 1 uL | 10X | 66 uL |
| **Hi-T4 DNA Ligase** | 500 U | 400 U/uL | 83 uL |
| **BsmBI-v2** | 15 U | 10 U/uL | 100 uL |
| **Water** | to 5 uL |  | 81 uL |
| **T4 Ligase Buffer** | 1 uL | 10X | 66 uL |
| **Hi-T4 DNA Ligase** | 500 U | 400 U/uL | 83 uL |
| **BsaI-HF-v2**<br>**or**<br>**BbsI-HF** | 15 U | 20 U/uL | 50 uL |
| **Water** | to 5 uL |  | 131 uL |

To set up reactions, combine 75 ng of the backbone part with 150 ng of each insert part in a PCR tube with 5 uL of the appropriate master mix and fill to 10 uL with water. Miniprepped parts give higher assembly efficiencies than linear PCR products.

| Golden Gate thermocycler protocol |
| --- |
| 20 min 37/42°C |
| 3 minutes 37/42°C |
| 4 minutes 16°C |
| Cycle 2-3 x30 |
| 10 minutes 50°C |
| 10 minutes 80°C |
| Hold 4°C |

**Supplementary Video 1: Example Acoustic Plate Reader scan**. (Right) The Acoustic Plate Reader is scanning six 96-well phantoms. (Left) The computer screen displays the real-time images of linear (left) and nonlinear (middle) contrast, as well as the Verasonics control interface (right).

**Supplementary Table 1: Oligos used for mutagenesis**. Sequences of the oligos that composed the four oligo pools used to create the GvpA/GvpB libraries. “Library Round” indicates the round of screening (first or second) in which the oligo was used, and “Sub-Library” indicates the pool in which it was synthesized.

**Supplementary Table 2: Custom-made MoClo parts**. Inventory of the MoClo parts added to the base EcoFlex system and used for cloning the constructs in this study.

**Supplementary Table 3: Ultrasound pulse sequences**. List of the parameters entered into the APR GUI to perform each scan in this study.

**Supplementary Table 4: PCR primers**. Sequences of the primers used to either amplify the oligo pools used to create the libraries, or to re-clone the best *gvpA/gvpB* mutants into Level 0 MoClo part vectors for assembly into expression constructs and subsequent validation.

## Notes

### Competing Interest Statement

The authors have declared no competing interest.

